# Development of a Reference Spatiotemporal Gait Data Set for Indian Subjects: A Pilot Study

**DOI:** 10.1101/080499

**Authors:** Raj Ramnani, Pooja Mukul, Siddhant Jain, Abhishek Tripathi

## Abstract

Interpretation of pathological gait has, for decades, offered insight into amputee anomalies and diseases such as cerebral palsy. Gait analysis has been actively used at the Bhagwan Mahaveer Viklang Sahatya Samiti for such purposes. The normative gait data used to compare data obtained from amputees, however, has been collected from laboratories under dissimilar conditions, skewing interpretation. Spatiotemporal gait parameters were extracted from 43 male Indian subjects using a 7.5m walkway and the *BTS Bioengineering GAITLAB* setup. Working under the hypothesis that a lack of cross-cultural validity was somewhat responsible for variations in normative gait, we attempted to develop a region-specific data set for use at the headquarters of the Jaipur Foot Organization Bhagwan Mahaveer Viklang Sahatya Samiti (BMVSS). Stratified random sampling was used to recruit subjects and measures were taken to ensure minimal effects of extraneous variables. Statistical analysis was performed on obtained data using one-way analysis of variance (ANOVA) to gauge the effect age and ethnicity had on normative values of the parameters investigated. We found statistically significant p-values for a few spatiotemporal parameters in the analysis of variance for both age and ethnicity. While the results were much less significant than initially hypothesized, the study proved an efficient way to create a normative gait data set exclusively for use at the gait laboratory of the Jaipur Foot Organization, thereby eliminating potential erroneous interpretation of pathological gait when comparing said gait to normative gait data obtained in laboratories under dissimilar conditions.

## 2. Introduction

Gait movement analysis is a widely used clinical procedure rendering rich, quantitative data, the applications of which include everything from orthopedic diagnosis facilitation to even biometric identification. Normative Gait data, however, is intrinsically flawed in that tremendous variations are prevalent amongst reported reference values of basic spatiotemporal parameters [1]. A primary cause of this variation, we hypothesized, might be the lack of cross-cultural validity in the existing reference data. Working under this assumption, and considering the need for region-specific data at Indian prosthetic clinics such as the Jaipur Foot Organization, where gait analysis facilitates prototype testing of prosthetic knees and other research, this pilot study aims to develop and analyze region-specific normative gait data using subjects of Indian origin exclusively (age: 10-59, N = 43). The applications of this data range from facilitation of better interpretation of pathological gait in Indian subjects to serving as a reference dataset for further research studies.

The essential purpose of the study is twofold—firstly, to ascertain whether or not a difference in ethnicity can affect normative gait, and if so, how significant is the intergroup variance caused, and secondly, to develop a normative gait dataset for Indian subjects.

## 3. Methods

### Subjects

43 healthy, male subjects within the age range of 10-59 were recruited as subjects using stratified random sampling in which non-amputated relatives of amputees seeking treatment/installment of a prosthetic at the Jaipur Foot Organization were randomly selected and invited to be a part of the study. All subjects were of Indian origin, and were born in and grew up in India. Subjects were also asked about which Indian state they and their family belonged to. Participants were made aware of their right to withdraw and the fact that their participation in the study was entirely voluntary; they could choose to leave at any point in time. A certificate of consent in a copy of the informed consent form was signed and dated by each subject, signifying their approval of the terms listed, and an additional copy was provided to them.

Prospective participants that presented with conditions known to affect gait, such as degenerative joint disease or disarticulation [2], were excluded. After exclusion, 43 male participants were left, all of which rated their health with descriptions ranging from normal to great, and gave informed consent before participation. Permission to approach relatives of amputees and approval for the study were granted by the director of research at the Jaipur Foot Organization, Dr. Pooja Mukul.

Since gait analysis requires subjects to be undressed, and most of the research team comprised males, women were not recruited given the strong social stigma prevalent and potential self-consciousness of women approached.

Apart from age, participants have been divided based on whether or not they were of Rajasthani origin. This was done in order to develop an additional, more region-specific dataset based off of the gait of the participants from Rajasthan, the Indian state where the Jaipur Foot Organization Headquarters and Research Center are based. This data set, we expected, would further improve interpretation of pathological gait in Rajasthani subjects, an obvious majority in the amputees that seek treatment at the Jaipur Foot Headquarters due to it’s location. Despite a majority of Rajasthani participants, a significant number of subjects and amputees, we observed, were also from Uttar Pradesh (UP), a state neighboring Rajasthan.

A comprehensive table details characteristics of participants below:

**Table 1:**
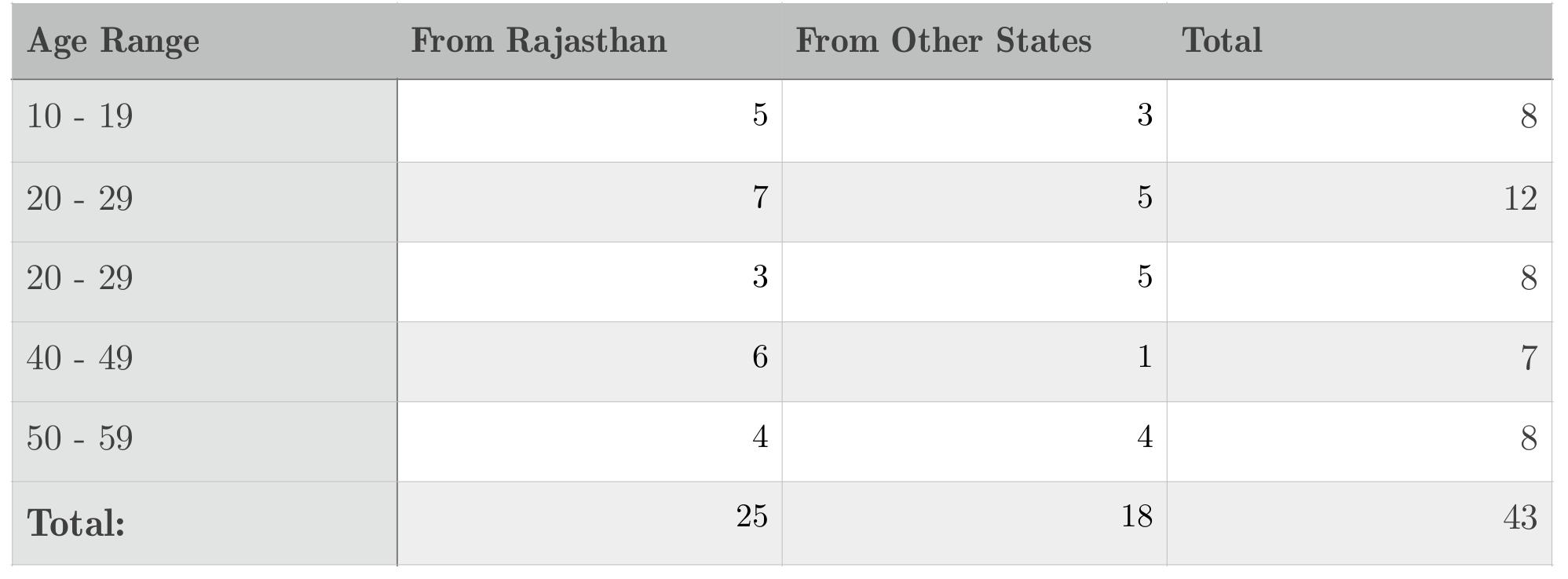
Participant Demographics

### Instrument Framework

Spatiotemporal gait measures were obtained using a BTS Bioengineering GAITLAB©, the setup of which was equipped with six infrared cameras, a force plate measuring kinetic data and a 7.5 meter walkway. [3] A computer was connected to the BTS Bioengineering Smart-DX Engine and ran the BTS Smart Clinic software that comprised an equipment connection diagnostic tool, a capturing tool and a cataloging application where sociodemographic data and anthropometric measures were collected and stored. Axes and wand calibrations were run multiple times to ensure optimal calibration and reliable data. Under the Simple Helen Hayes protocol used, BTS infrared markers were attached to the bodies of the subjects prior to the walks in fifteen distinct physiological locations below the anatomical transverse plane for the infrared cameras to detect. The fifteen markers are shown and labeled alongside [4]:

**Figure 1:**
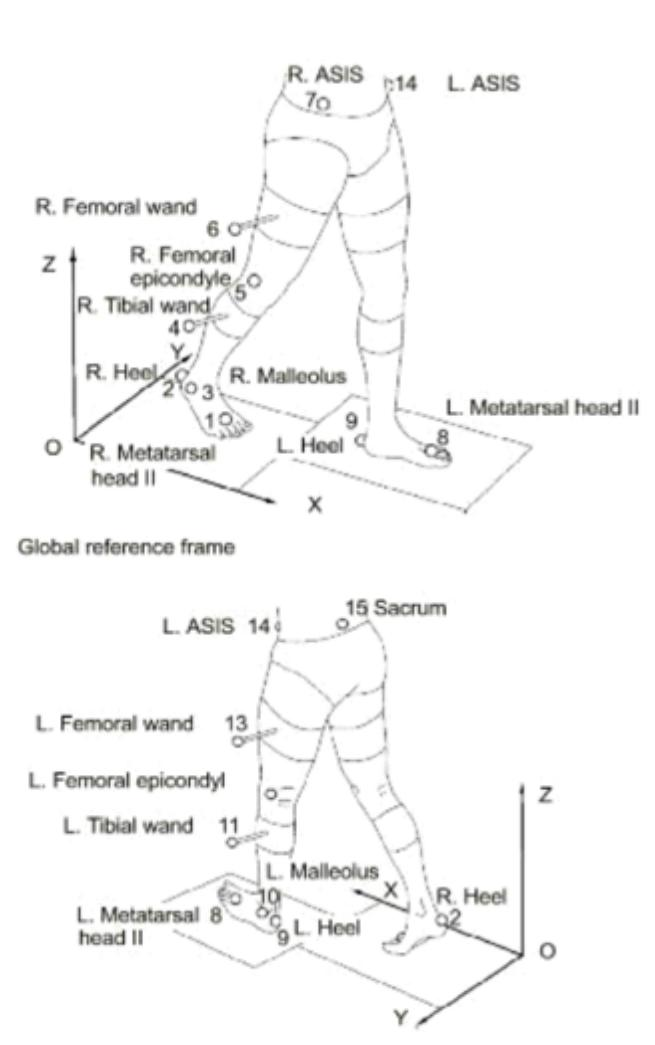
Marker Placement Anatomical Locations [4]

### Procedure

After head and wand infrared markers were attached using double adhesive tape and belt-like bands respectively, subjects walked across the walkway twice in their undergarments, once towards each end. Keeping the objective of simulating realistic gait and retaining ecological validity in mind, subjects were asked to disregard the laboratory setting and the markers when walking, and to walk how they normally did in their daily lives [5]. Measures were taken to ensure that they began and ended their walk a good three feet (~1m) before and after the walkway in order to minimize the effect of acceleration as an extraneous variable. Instead of running one trial wherein a subject would turn at the end of the walkway and walk the other way, two separate trials were run so as to avoid effects a reduction in velocity and step length during the turn would have had on the variance and mean of the data collected in each frame set capturing a complete gait cycle [6].

The values of 13 different distance-temporal gait parameters were collected for each subject. A three-axis force plate was in place under the mat on the walkway [7]. Some parameters, such as stance phase and swing phase, were quantified in two different mediums (% of gait cycle, or %GC, and seconds, or s.), and thereby treated as disparate variables, each offering unique insight into normative gait [8].

The gait parameters used in this study are fundamental kinematic measures detailed below:

**Table 2:** Parameters Investigated

| Temporal Parameters | Spatial Parameters |
| --- | --- |
| Stance Phase (%) | Step Length (m) |
| Swing Phase (%) | Velocity (m/s) |
| Double Support (%) | Swing Velocity (m/s) |
| Stance Phase (s) | Stride Length (m) |
| Swing Phase (s) | Step Width (m) |
| Stride Time (s) | Mean Velocity (m/s) |
| Cadence (step/min) |  |

Operational Definitions of all aforementioned parameters can be found in the tables below:

**Table 3:** Operational Definitions

| Temporal Parameters | Operational Definitions |
| --- | --- |
| Stance Phase (%) | Stance Time normalized to Stride Time |
| Swing Phase (%) | Swing Time normalized to Stride Time |
| Double Support (%) | Double Support Time normalized to Stride Time |
| Stance Phase (s) | Time elapsed between final and initial contact of a singel footfall |
| Swing Phase (s) | Time elapsed between final contact of one footfall and initial contact of the next footfall of the same foot |
| Stride Time (s) | Time elapsed between initial contacts of consecutive footfalls of the same foot |
| Cadence (step/min) | Step rate - Number of steps taken per minute. |

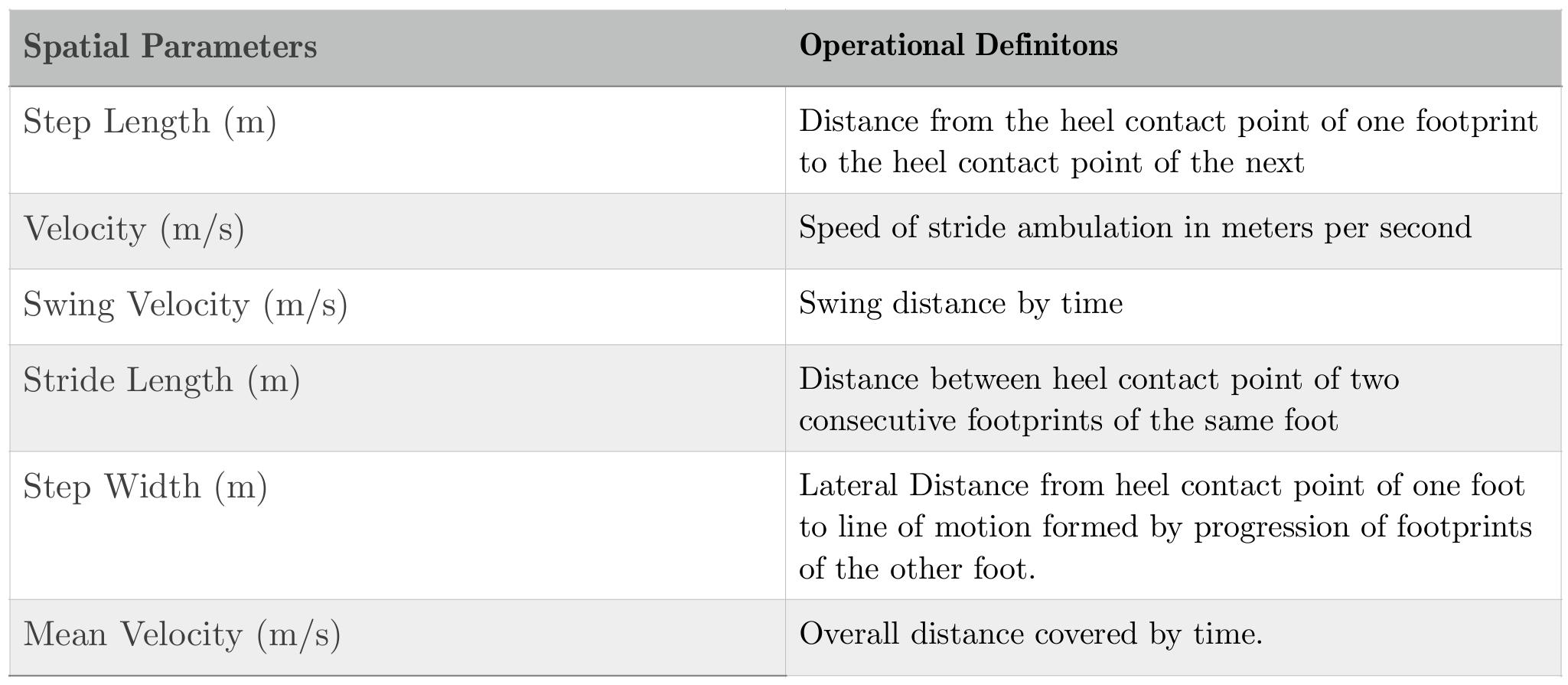

| Spatial Parameters | Operational Definitons |
| --- | --- |
| Step Length (m) | Distance from the heel contact point of one footprint to the heel contact point of the next |
| Velocity (m/s) | Speed of stride ambulation in meters per second |
| Swing Velocity (m/s) | Swing distance by time |
| Stride Length (m) | Distance between heel contact point of two consecutive footprints of the same foot |
| Step Width (m) | Lateral Distance from heel contact point of one foot to line of motion formed by progression of footprints of the other foot. |
| Mean Velocity (m/s) | Overall distance covered by time. |

A clinical gait report, created especially for gait analysis at the Jaipur Foot Organization, was generated after tracking and processing trial data with the Smart Clinic software, and relevant kinematic data was extracted from said report.

### Data Analysis

The *IBM SPSS Desktop 24.0* was used for data collection, processing, and statistical analysis.

## 4. Results

In order to compare the means of spatiotemporal gait parameters between Caucasian and Indian subjects, we carried out a one-way analysis of variance (ANOVA) test [9], and, in essence, partitioned variability into that caused by difference in ethnicity and that caused by other factors, such as age (or variability within the values obtained for subjects of a particular ethnic group). The results obtained are tabulated below:

### One-Way ANOVA—Completely Randomized

**Table 4:**
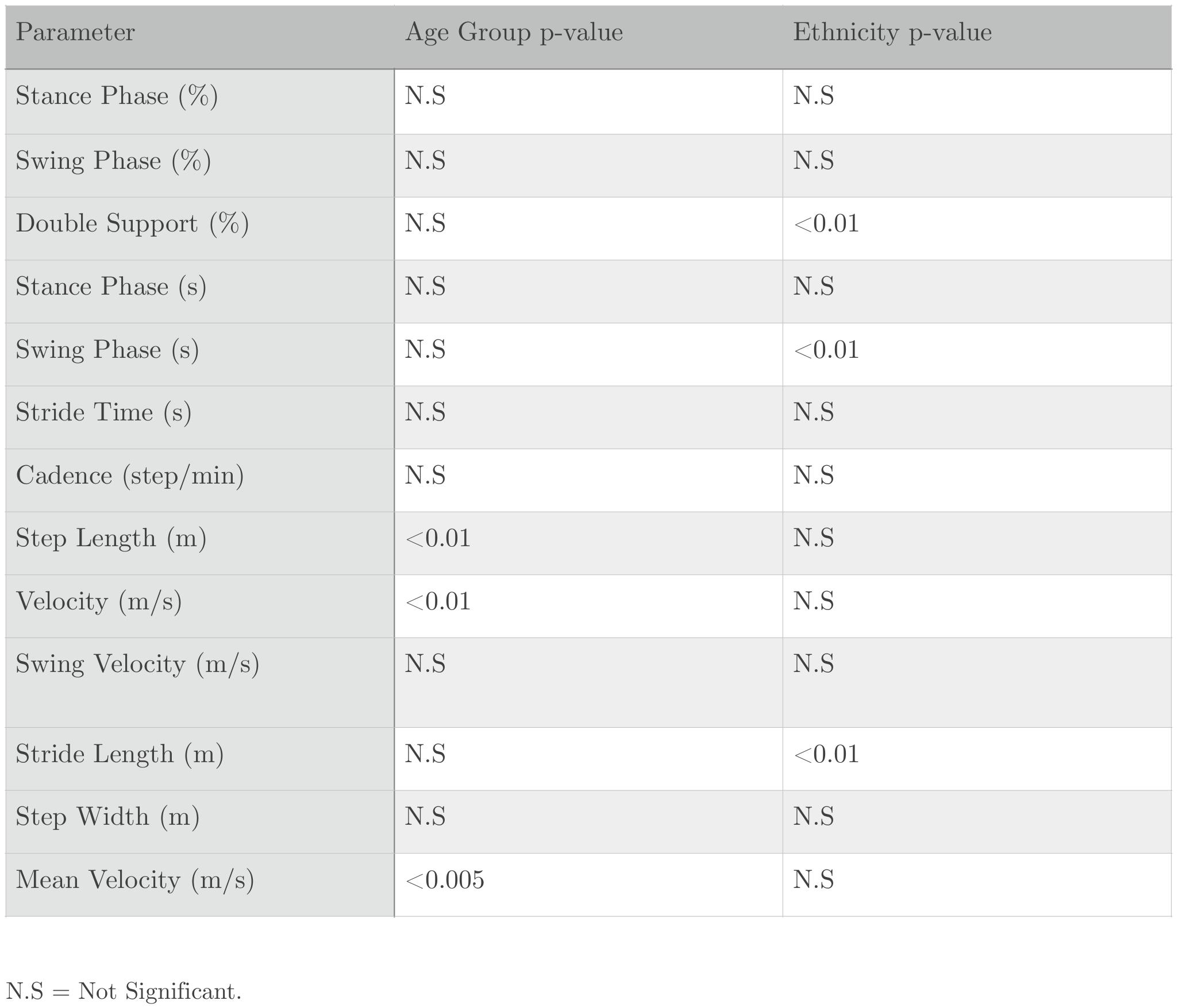
ANOVA p-values

When looking at ethnicity, we found statistically significant p-values for Double Support, Swing Phase, and Stride Length, but other parameters rendered p-value above 0.01 in varying degrees.

When looking at age as a factor influencing normative values, we discovered that statistical significance was only prevalent in Step Length, Velocity, and Mean Velocity.

Clearly, lab conditions, age, and a lack of cross-cultural validity affect reference values of certain gait parameters, albeit a few. The results signified that while the sample was too small to make conclusive interpretations, it can be inferred that such statistical significance would most likely be observed within a larger and more representative sample size as well. The pilot study served its purpose by illustrating a correlation, and therefore, similar procedures will be carried out with a larger sample to effectively establish a cause and effect relationship.

Descriptive statistics detailing means, standard deviations, and standard error for each parameter can be found below:

**Table 5:** Descriptive Statistics

| Parameter | Type | N | Mean | Std. Deviation | Std. Error |
| --- | --- | --- | --- | --- | --- |
| Stance Phase | R(%) | 43 | 58.8 | 4.85904 | ~1.3 |
| Stance Phase | L(%) | 43 | 58.2 | 5.81899 | ~1.7 |
| Swing Phase | R(%) | 43 | 41.2 | 4.44032 | ~1.3 |
| Swing Phase | L(%) | 43 | 41.8 | 7.28426 | ~1.7 |
| Double Support | R(%) | 43 | 11.2 | 2.02679 | ~1.5 |
| Double Support | L(%) | 43 | 12.4 | 1.92245 | ~1.5 |
| Stance Phase | R(s) | 43 | 0.56 | 0.05561 | ~.04 |
| Stance Phase | L(s) | 43 | 0.56 | 0.06822 | ~.03 |
| Swing Phase | R(s) | 43 | 0.46 | 0.04810 | ~.03 |
| Swing Phase | L(s) | 43 | 0.46 | 0.08040 | ~.03 |
| Stride Time | R(s) | 43 | 1.02 | 0.11046 | ~.02 |
| Stride Time | L(s) | 43 | 1.06 | 0.07234 | ~.03 |
| Cadence | (step/min) | 43 | 114.593 | 6.56892 | ~3.9 |
| Step Length | R(m) | 43 | 0.66 | 0.14126 | ~.04 |
| Step Length | L(m) | 43 | 0.64 | 0.15111 | ~.03 |
| Velocity | R(m/s) | 43 | 1.31 | 0.14198 | ~.05 |
| Velocity | L(m/s) | 43 | 1.29 | 0.14538 | ~.06 |
| Swing Velocity | R(m/s) | 43 | 3.19 | 0.28295 | ~.22 |
| Swing Velocity | L(m/s) | 43 | 3.05 | 0.36752 | ~.19 |
| Stride Length | R(m) | 43 | 1.38 | 0.23308 | ~.06 |
| Stride Length | L(m) | 43 | 1.36 | 0.12374 | ~.07 |
| Step Width | R(m) | 43 | 0.14 | 0.06204 | ~.02 |
| Step Width | L(m) | 43 | 0.17 | 0.06204 | ~.02 |
| Mean Velocity | (m/s) | 43 | 1.27 | 0.14000 | ~.05 |

A table of the normative values can be found below:

**Table 6:** Normative Values

| Parameters | Right | Left |
| --- | --- | --- |
| Stance Phase (%) | 58.8±1.3 | 58.2±1.7 |
| Swing Phase (%) | 41.2±1.3 | 41.8±1.7 |
| Double Support (%) | 11.2±1.5 | 12.4±1.5 |
| Stance Phase (s) | 0.56±.04 | 0.56±.03 |
| Swing Phase (s) | 0.46±.03 | 0.46±.03 |
| Stride Time (s) | 1.02±.02 | 1.06±.03 |
| Cadence (step/min) | 114.593±3.872 | 114.593±3.872 |
| Step Length (m) | 0.66±.04 | 0.64±.03 |
| Velocity (m/s) | 1.31±.06 | 1.29±.05 |
| Swing Velocity (m/s) | 3.19±.22 | 3.05±.19 |
| Stride Length (m) | 1.38±.06 | 1.36.07 |
| Step Width (m) | 0.14±.02 | 0.17±.02 |
| Mean Velocity (m/s) | 1.27±.05 | 1.27±.05 |

## 5. Discussion

While literature reveals that a larger sample size (N>200) is more representative of the target population [10], our study provides a prototype-like, region-specific dataset (Appendix 2) that can be expected to marginally improve accuracy of interpretation of pathological gait tested in similar conditions. (The dataset details normal parameter values and can be found in Appendix 2).

Statistical analysis using one-way analysis of variance proved that there was in fact a statistically significant variation in a few normative gait parameter values caused by differences in age and ethnicity, namely double support, swing phase, cadence, step length, velocity, and mean velocity. Although such significance was not observed in most parameters, it’s safe to say that individual differences leading to variations in normative gait can be prompted by factors such as age and ethnicity. [11]

We were able to create a reference data set for use exclusively at the Jaipur Foot Organization so as to avoid erroneous interpretation of pathological gait that may arise when comparing said gait to normative data obtained elsewhere in dissimilar conditions.

### Concerns regarding ecological validity

The measurements and results of the project were dependent on test conditions, circumstances that can’t be emulated exactly. In ordinary lab conditions, however, results should approximately round off to similar figures, and so our data is truly only suitable for comparison to data obtained under standard laboratory conditions on a short walkway. As demonstrated by Öberg et al, gait analysis data should only be comprehended with accurately defined test conditions. [12]

It is also crucial to note that all subjects walked at self-selected speeds, which may—to some extent—have led to a lack of external validity.

These observations were more or less supported by the results of studies by Öberg et al and Dahlstedt et al [13], although the mean for self-selected gait speed was negligibly higher to that found by Öberg et al, who used a short laboratory walkway. Our mean gait speed of ~127 cm/s was also marginally lower than that found by Murray et al [14](151cm/s for men), who sampled free gait outdoors on a long walkway. Clearly, there is distinct difference between gait indoors and outdoors and between gait sampled on a short walkway and a long one—a distinction that calls for separate reference datasets for each test condition.

### Variations caused by age

While aging appeared to be contributing to a reduction in step length and velocity, there was only a negligible change in cadence (step frequency). In our study, we found statistically significant gait differences related to age and observed an average decrease of 0.3 %/year in velocity. Ordinarily, this number ranges from 0.1% to 0.7%, [15] differences within which arise primarily due to a discrepancy in test conditions.

### Variations caused by ethnicity

Ethnicity was observed to prompt variations in step length in addition to double support and swing phase. While ethnicity wasn’t as strongly responsible as we initially hypothesized, our data could potentially improve gait interpretation of amputees analyzed at the Jaipur Foot Organization and build the foundation for in-depth studies with a larger sample more representative of the population.

It is also important to note that being brought up and having lived in an Indian sociocultural context are factors that contributed to the aforementioned variance in addition to ethnicity.

### Future Scope

Our study is an extension of and supports Majernik’s guideline for individual laboratories to develop a reference data set of their own, and maximize that objectivity of gait analysis by not relying on databases generated at other laboratories even if they are equipped with similar motion analysis systems [16]. While the current undertaking will only benefit the gait and movement analysis laboratory at the Jaipur Foot Organization, a collaboration to expand on this and eventually create a more robust standard database is underway. Machine Learning algorithms are looking to be used to allow labs to create their own databases with ease (based on a few test subjects) to facilitate better interpretation of pathological gait. This dataset will serve as a test sample in the bigger project.

## 6. Acknowledgements

We would like to thank our school, our academic counsellor, Mr. Ayush Periwal, our families, and our friends for their unconditional support throughout our undertaking of this endeavor.

We are grateful to MIT OpenCourseWare for their online course on Developing World Prosthetics and to Duke University for their instructional videos on “Statistics with R” on Coursera.

Above all else, Dr. Pooja Mukul’s (Director of Research at BMVSS) help and guidance were indispensable and made this project possible. We are also indebted to Mr. D.R. Mehta for offering us a research internship at the Bhagwan Mahaveer Viklang Sahatya Samiti/Jaipur Foot Organization.

Lastly, we would also like to express our gratitude to the rest of the team at the Paul Hamlyn Research Facility at the Jaipur Foot Organization that helped us out with our project and let us use their Gait and Movement Analysis Lab.

